# Neuronal expression of human amyloid-β and Tau drives global phenotypic and multi-omic changes in *C. elegans*

**DOI:** 10.1101/2023.06.01.542377

**Authors:** Angelina Holcom, Matias Fuentealba, Renuka Sivapatham, Christina D. King, Hadley Osman, Anna Foulger, Dipa Bhaumik, Birgit Schilling, David Furman, Julie K. Andersen, Gordon J. Lithgow

## Abstract

Alzheimer’s disease (AD) and Alzheimer’s related diseases (ADRD) are prevalent age-related neurodegenerative disorders characterized by the accumulation of amyloid-β (Aβ) plaques and Tau neurofibrillary tangles. The nematode *Caenorhabditis elegan*s (*C. elegans*) serves as an invaluable model organism in diseases of old age-due to its rapid aging. Here we performed an unbiased systems analysis of a *C. elegans* strain expressing both Aβ and Tau proteins within neurons. We set out to determine if there was a phenotypic interaction between Aβ and Tau. In addition, we were interested in determining the temporal order of the phenotypic and multi-omic (geromic) outcomes. At an early stage of adulthood, we observed reproductive impairments and mitochondrial dysfunction consistent with disruptions in mRNA transcript abundance, protein solubility, and metabolite levels. Notably, the expression of these neurotoxic proteins exhibited a synergistic effect, leading to accelerated aging. Our findings shed light on the close relationship between normal aging and ADRD. Specifically, we demonstrate alterations to metabolic functions preceding age-related neurotoxicity, offering a resource for the development of new therapeutic strategies.

## Main

Alzheimer’s disease (AD) and Alzheimer’s related diseases (ADRD) represent the most prevalent forms of dementia, affecting millions of individuals worldwide^1^. The greatest risk factor for ADRD is aging. Pathological hallmarks of these disorders include the formation of amyloid-β (Aβ) plaques, neurofibrillary tangles composed of hyperphosphorylated Tau protein, glial cell activation, and impairment of the blood-brain barrier.

The interaction between Aβ and Tau in AD has been closely linked to hypometabolism in the posterior cingulate cortex, which precedes memory decline^2^. Aβ presence accelerates Tau oligomerization, hyperphosphorylation, and propagation^2–4^. In mouse models, co-expression of Aβ and Tau leads to early synapse loss and synergistic effects on defects in mitochondrial function and turnover^2^. Patients with both Aβ and Tau, as determined by cerebrospinal fluid (CSF) analysis and MRI/PET methods, experience an amplified rate of cognitive decline compared to individuals with only Aβ, only Tau, or neither^5^.

Extensive research has focused on finding disease-modifying treatments for AD, including inhibition of Aβ production by targeting the cleavage of APP by β and γ secretase, along with immunotherapy, and inhibitors of Aβ aggregation. However, these approaches have not succeeded in halting disease progression^6^. Consequently, there is need to further explore therapeutic targets, such as targeting multiple factors of the disease concurrently. A potential target could be centered around interactions between Aβ and Tau, instead of focusing on the individual proteins associated with the disease process.

We investigated the multi-omic (geromic) consequences of pan-neuronal, simultaneous expression of human Aβ and Tau in the model organism *C. elegans* leveraging its short 20-day lifespan. In previous studies transgenic *C. elegans* models have expressed human Aβ in the body muscle or neurons, resulting in a range of phenotypes^7–9^.

Similarly, investigations of Tau expression in *C. elegans* have focused on expression of human Tau containing mutations associated with frontotemporal dementia (FTD), a disorder related to AD^10^. Recently, there have been two models that have combined human Aβ and Tau into a single *C. elegans* model and observed decreased lifespan and progeny viability along with increased neuron loss, motor dysfunction and protein aggregation^11,12^. In addition, there were alterations on a transcriptomic level including changes in UDP-glucuronosyltransferase, protein phosphorylation, and aging genes^11^. These models all express the human form of the toxic proteins due to them not being endogenously found within *C. elegans*. Despite these proteins not being endogenously expressed these transgenic models have each successfully reproduced certain aspects of AD-like phenotypes, including insoluble Tau accumulation, age-related uncoordinated movement, and neurodegeneration^10^.

To examine the impact of simultaneous expression of Aβ and Tau, we crossed *C. elegans* strains expressing pan-neuronal Aβ and pan-neuronal 4R1N Tau with V337M mutation, which allows us to further examine the interaction of the proteins but with the addition of the V337M mutation to accelerate Tau aggregation^13,14^. This model allowed us to explore the impact of these AD-related proteins on whole-organism protein insolubility, transcriptome dynamics, and metabolomics across different stages of life, including young, middle, and old age. Significantly, this model allowed us to uncover insights into potential early-life effects on reproductive functions and the intricate relationship between normal aging and the development of chronic age-related pathologies.

## Results

### Systemic phenotypes observed with expression of Tau or Aβ

To investigate the impact of expression of WT human Aβ and 4R1N Tau containing the V337M mutation on protein aggregation^13,14^, we generated a *C. elegans* strain with simultaneous pan-neuronal expression of both proteins (Fig. 1A). This was achieved by crossing the existing pan-neuronal Aβ strain, CL2355, with the pan-neuronal Tau strain, CK10. The CL2355 strain uses the *snb-1* promoter to drive expression of Aβ in the nerve ring, ventral cord, dorsal cord and neuronal cell bodies^15^. While the CK10 strain uses the *aex-3* promoter to drive expression of Tau in nearly all the neurons including the major ganglia next to the nerve ring, ventral cord and tail ganglia^16^. This newly generated strain was first used to determine whether the expression of Aβ and Tau leads to enhanced phenotypic dysfunction. We initially focused on testing for a phenotype seen in AD, alteration in mitochondrial function. We tested mitochondrial function by measuring the worm’s ability to move and the rate of oxygen consumption of the mitochondria. Movement patterns of the worms were quantitated by measuring the number of body bends (thrashing rate) in a given time. The Tau;Aβ strain exhibited a significantly decreased thrashing rate compared to the WT, Tau, and Aβ strains (Fig. 1B). Mitochondrial function was quantitated by measuring the rate of oxygen consumption using a Seahorse XFe96 Analyzer. The Tau;Aβ strain displayed increased average oxygen consumption rates (OCR) compared to the Aβ strain while the Tau strain had increased average OCR compared to WT and Aβ strains, suggesting heightened mitochondrial activity (Fig. 1C). These findings imply that the expression of Tau and Aβ have a synergistic effect on the body movement phenotype while Tau appears to have a stronger effect on mitochondrial OCR.

**Figure 1:**
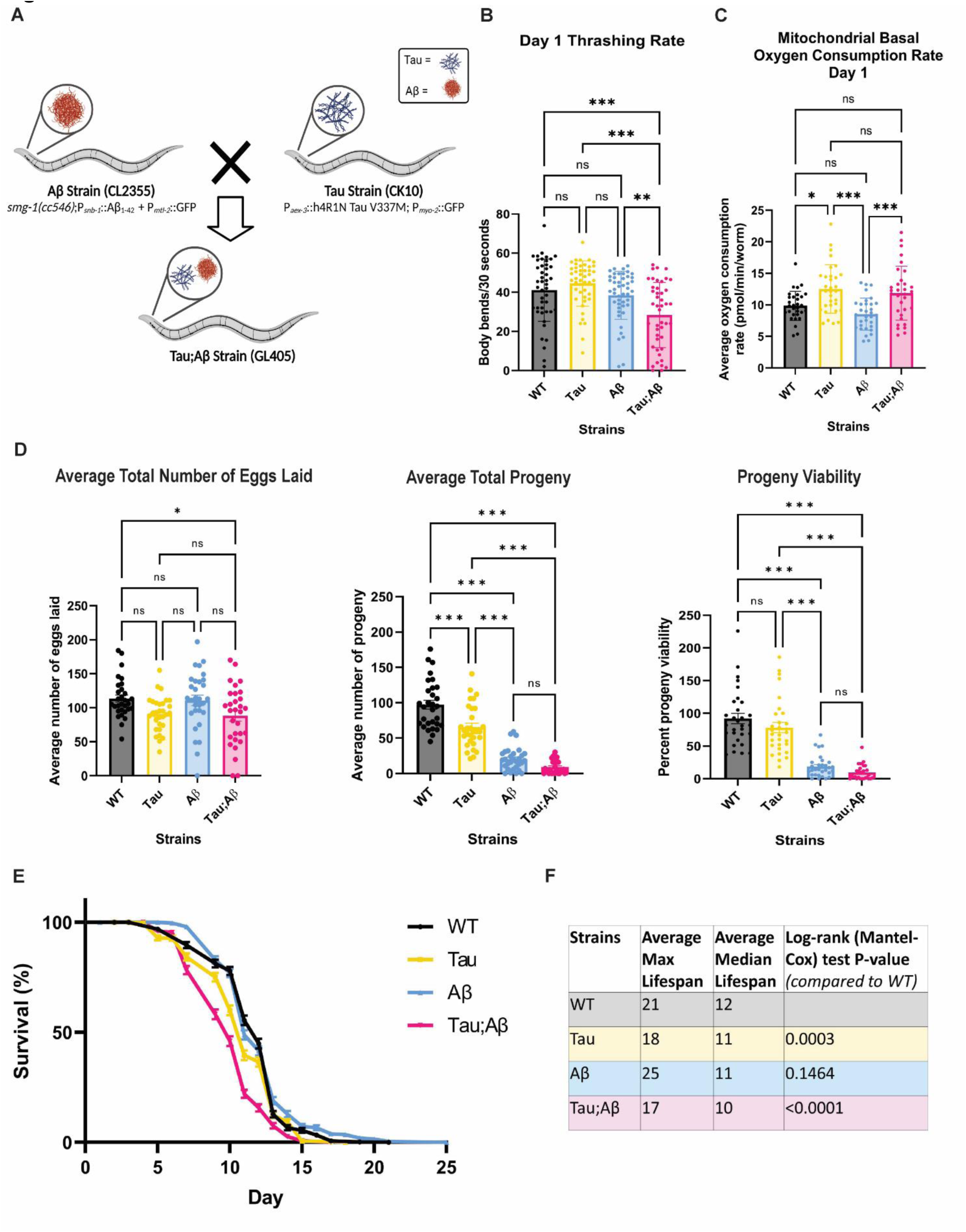
Systemic phenotypes observed in the context of expression of Tau and Aβ. (A) Diagram of the cross used to create the dual model simultaneously expressing both human WT Aβ and mutant Tau. The newly generated strain expresses pan-neuronal human WT Aβ and 4R Tau containing the FTDP-17 mutation V337M as well as both pharyngeal and intestinal GFP. (B) Average thrashing rates of the 4 strains on day 1 of adulthood. 3 biological replicates with 10-20 worms per replicate. Error bars represent mean ± SEM. (C) Average basal mitochondrial oxygen consumption rate (OCR) of day 1 adult WT, Tau, Aβ and Tau;Aβ strains. 3 biological replicates with 8 technical replicates, which had around 20 worms in each technical replicate. Error bars represent mean ± SEM. (D) The average total number of eggs laid in the 4 strains with 3 biological replicates, 10 worms per replicate. Error bars represent mean ± SEM. The total progeny in the 4 strains with 3 biological replicates, 10 worms per replicate.. Error bars represent mean ± SEM. The percent of progeny viable was determined by the total number of progeny hatched and total eggs laid. Ordinary one-way ANOVA with Tukey’s multiple comparisons test. Error bars represent mean ± SEM. (E) Average lifespan for WT, Tau, Aβ and Tau;Aβ strains. Includes table of average max and median lifespan and log-rank test p-value of AD strains compared to WT. 3 biological replicates with 3 technical replicates that had 40-60 worms each. Error bars represent mean ± SEM.

In the maintenance of the Tau;Aβ strain, we observed altered hermaphrodite self-fertility. This observation drove us to characterize this possible toxicity phenotype. The number of eggs laid (fecundity) and the number of eggs that hatched (an early indication of fertility) were measured until the exhaustion of egg laying. The Tau and Tau;Aβ strains exhibited a decreased average total number of eggs laid, while the Aβ and Tau;Aβ strains showed a significantly decreased total progeny (Fig. 1D).

As a measure of overall health, we performed a survival assay on adult hermaphrodites, revealing that the Tau;Aβ expressing strain had a decreased median and maximum lifespan compared to the other three strains (Fig. 1E&F). This may represent accelerated aging.

These results indicate that the pan-neuronal expression of Aβ and Tau exerts systemic effects on the worms, with Aβ affecting progeny production, Tau influencing mitochondrial OCR, and the combined expression of both proteins leading to reduced body movement and lifespan. These observations underscore the impact of Tau and Aβ interactions on the phenotypic characteristics of the worms.

### Synergistic effects of Aβ and Tau expression on metabolic profiles in young animals

To study the systemic consequences of expressing human neurotoxic proteins in *C. elegans*, we conducted global metabolomic analysis of the WT, Tau, Aβ, and Tau;Aβ expressing strains on day 1, 3, 5 and 8 of worm adulthood. At day 1 of adulthood the Tau and Aβ strains exhibited relatively minor differences in metabolite profiles compared to the WT strain (Fig. 2A). However, the Tau;Aβ expressing strain displayed extensive alterations in metabolite profiles compared to the WT strain (Fig. 2A), with the Aβ expressing strain resembling this trend based on correlation analysis (Fig. 2B).

**Figure 2:**
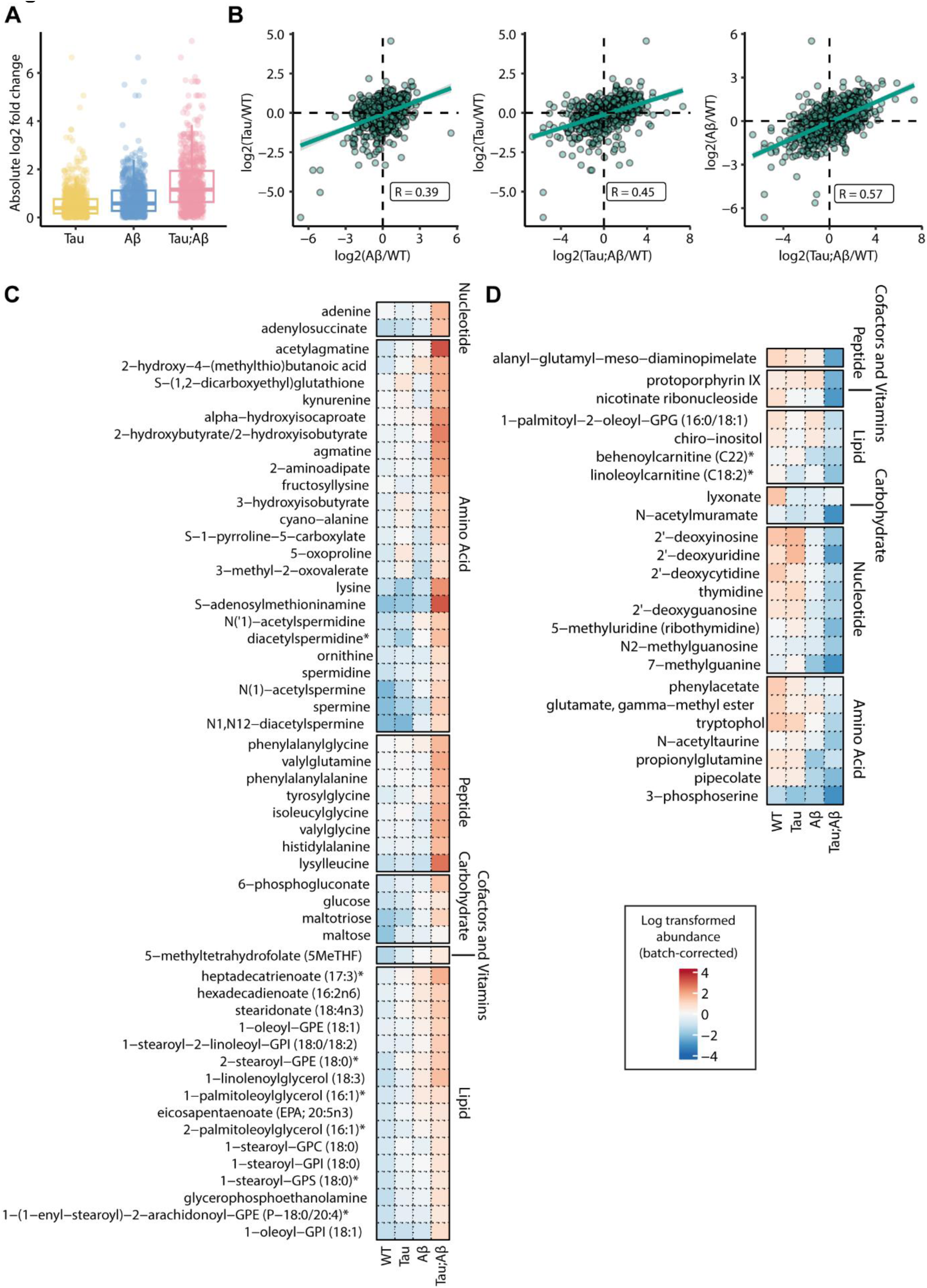
Expression of Tau and Aβ synergistically alters metabolite levels. (A) Bar graph of fold metabolite change in day 1 Tau, Aβ and Tau;Aβ compared to WT. (B) Pearson’s product-moment correlation of fold metabolite change between Aβ vs WT and Tau vs WT, and Tau;Aβ vs WT. (C) Heatmap of metabolites and their super pathways, which are significantly increased in Tau;Aβ compared to WT, Tau and Aβ. (D) Heatmap of metabolite and their super pathways, which are significantly decreased in Tau;Aβ compared to WT, Tau and Aβ.

Further examination of the metabolites altered in the day 1 Tau;Aβ expressing strain revealed contrasting abundance compared to the WT strain (Fig. 2C). The metabolites significantly increased in the Tau;Aβ strain primarily belonged to the categories of lipids and amino acids, specifically metabolites involved in spermidine and spermine biosynthesis, biotin metabolism and butyrate metabolism (Fig. 2C). In contrast the metabolites significantly decreased were in the categories of nucleotides and amino acids involved in folate metabolism and glycine and serine metabolite(Fig. 2C). These changes could connect back to the mitochondrial phenotype as we see changes in mitochondrial-related metabolites including spermidine and spermine biosynthesis along with glycine and serine metabolism.

In summary, our metabolomic analysis highlighted the synergistic effects of Aβ and Tau expression on the metabolic profiles in young adult animals. The Tau;Aβ expressing strain exhibited substantial alterations in metabolites, particularly in lipid and amino acid metabolism.

### Expression of Tau and Aβ disrupts metabolism throughout life

To gain deeper insights into the metabolic changes associated with aging in the WT, Tau, Aβ, and Tau;Aβ strains, we examined day 3, 5, and 8 metabolomic data. When looking at the number of metabolites that significantly change on day 3, 5 or 8 when compared to day 1 we observed that significant changes were primarily evident on day 8 for the WT and Tau expressing strains (Fig. 3A). The Aβ expressing strain exhibited a slight increase in significantly altered metabolites on day 5, which became more prominent on day 8. In contrast, the Tau;Aβ expressing strain showed the most significant changes on all three days, indicating a potential additive and/or synergistic relationship resulting from the simultaneous expression of both proteins in neurons.

**Figure 3:**
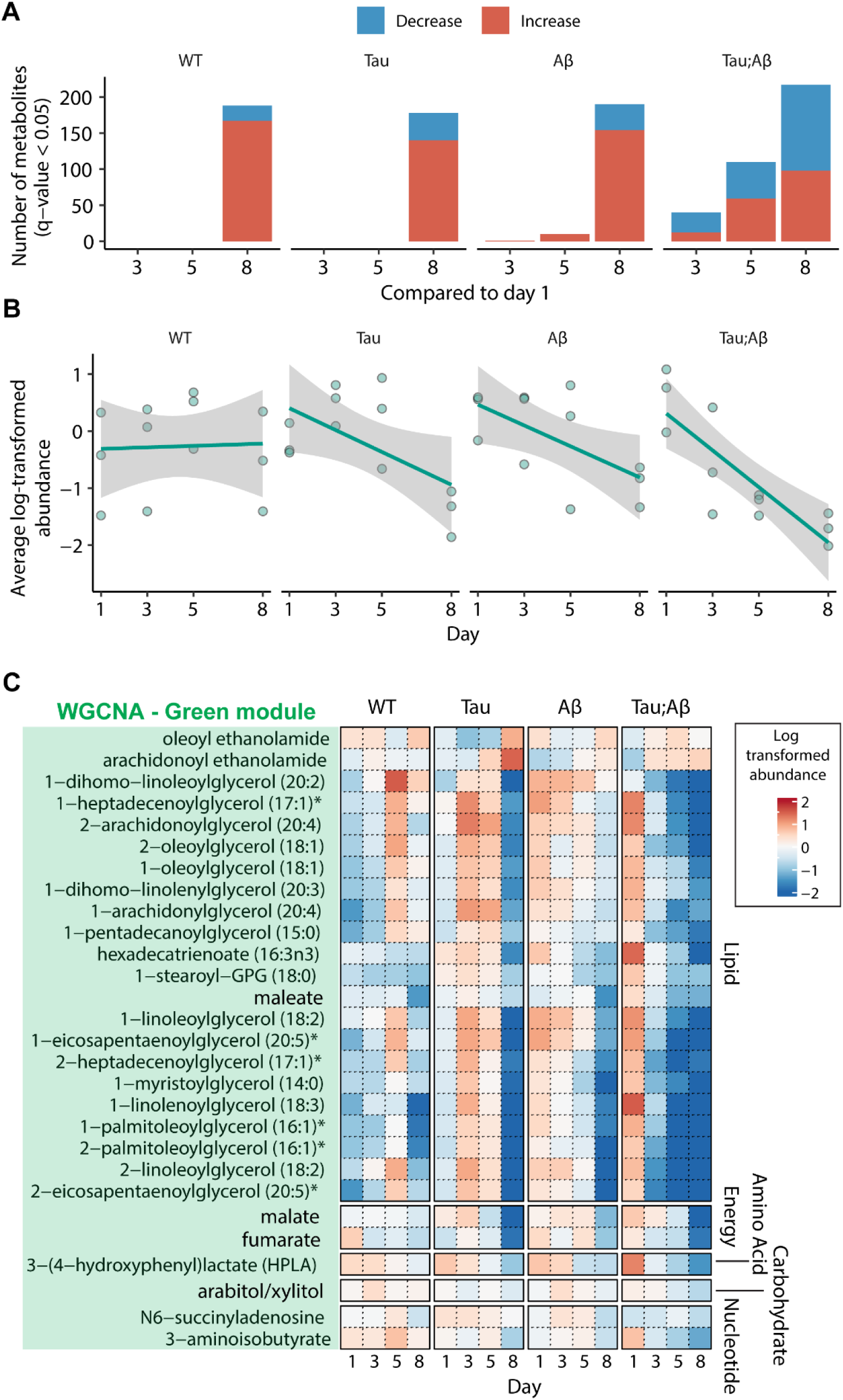
Expression of Tau and Aβ enhances metabolic changes with age. (A) Bar graphs of the number of metabolites significantly changing in day 3, day 5 and day 8 vs day 1 in all 4 strains. (B) Graph of the WGCNA green module with all 4 strains from day 1 to day 8 with 3 biological replicates. The y-axis value represents the eigengene value, which is the summarized abundance of the metabolites in the green module. (C) Heatmap of metabolites and their super pathways in the green module for all strains on day 1, 3, 5 and 8.

Comparing the metabolite levels on day 8 compared to day 1 for each strain, we identified an overlap in metabolites altered in abundance on day 8 across all four strains. However, the Tau;Aβ strain had a greater number of unique lower abundance metabolites that did not overlap with the other strains (Supplementary Fig. 1A&B).

When comparing the AD strains to the WT on day 3, 5, and 8, the Tau;Aβ strain exhibited the most significant alterations in metabolites on day 5 (Supplementary Fig. 1C). This was also visualized in Supplementary Figs. 1D, E and F where the log fold change of the metabolites in AD strains vs WT on the different days are depicted. The Tau;Aβ strain were shown to have a slight increase in log fold change on day 3 and a larger change on day 5 in comparison to the other AD strains. On day 8 the AD strains appear to have an increase in metabolite fold change with the Tau;Aβ strain still having the largest change (Supplementary Fig. 1F).

We performed a weighted correlation network analysis (WGCNA) to cluster metaboliets with similar abundance profiles into different modules (Supplementary Fig. 2A&B). One of the modules, labeled “green,” displayed relatively consistent levels of clustered metabolites in the WT strain with age (Fig. 3B). The Aβ, Tau and Tau;Aβ strains showed elevated metabolite abundance levels on day 1 and decreasing abundance from day 3 to day 8. The Tau;Aβ strain exhibited a synergistic effect with a more significant decrease in metabolite abundance with age compared to Aβ and Tau (Fig. 3B). This trend can also be seen in the heatmap of the green module metabolites (Fig. 3C). The metabolites in this green module consist mostly of lipids, such as monoacylglycerols which are glycerolipids that are found to be elevated in AD brains and alterations are found to occur in early stages of the disease^17^.

In addition to the “green” WGCNA module, we also identified additional modules labeled “salmon”, “pink”, and “grey60”. The “salmon” module represents a cluster of metabolites that are elevated in all ages in the Tau;Aβ strain, while they increased with age in the WT, Tau and Aβ strains (Supplementary Fig.2C). These metabolites are associated with glycerophospholipid metabolism and purine metabolism (Supplementary Fig.2D). The “pink” module represents a cluster of metabolites that are elevated on day 1 in the Tau;Aβ strain and decreased with age, while having constant levels in the Tau and Aβ strains. (Supplementary Fig.2E). These metabolites are associated with biosynthesis of unsaturated fatty acids (Supplementary Fig.2F). The “grey60” module represents a cluster of metabolites that are elevated on day 1 in the Tau;Aβ strain and decreased with age, while in the WT, Aβ and Tau strains the metabolites are decreased on day 1 and increased with age (Supplementary Fig.2G). These metabolites are associated with glycolysis/gluconeogenesis and pentose phosphate pathway (Supplementary Fig.2H).

Elevated pathways in the Tau;Aβ strain, including energy and fat, suggests that the expression of Tau and Aβ leads to disruptions in energy metabolism, lipid metabolism, and mitochondrial function, which may contribute to the observed changes in the mitochondrial, body movement, fertility and fecundity phenotypes associated with the Tau;Aβ strain.

### Expression of Aβ and Tau drives insoluble protein accumulation in young animals

To further study the systemic effects of the expression of Aβ and Tau, we were interested in determining if their expression leads to the aggregation of other proteins as we had previously observed in *C. elegans* with aging, via induction by other stressors, such as iron^18–20^. We collected insoluble proteins from Day 1 adult worms of WT, Tau, Aβ, and Tau;Aβ strains, followed by mass spectrometric analysis (see Supplementary Table 1-4) using data-independent acquisition^21,22^.

A comparison of the insoluble proteins in Tau vs WT, Aβ vs WT, and Tau;Aβ vs WT strains demonstrated an overlap among all three AD strains, with the Aβ expression alone significantly increased the abundance of insoluble proteins compared to WT worms. The Tau;Aβ expressing strain showed the most substantial alterations similar to the metabolomics data, where we saw changes in serine, glycine metabolism which plays a role in protein synthesis which may have a role in this proteomic finding (Fig.4A).

**Figure 4:**
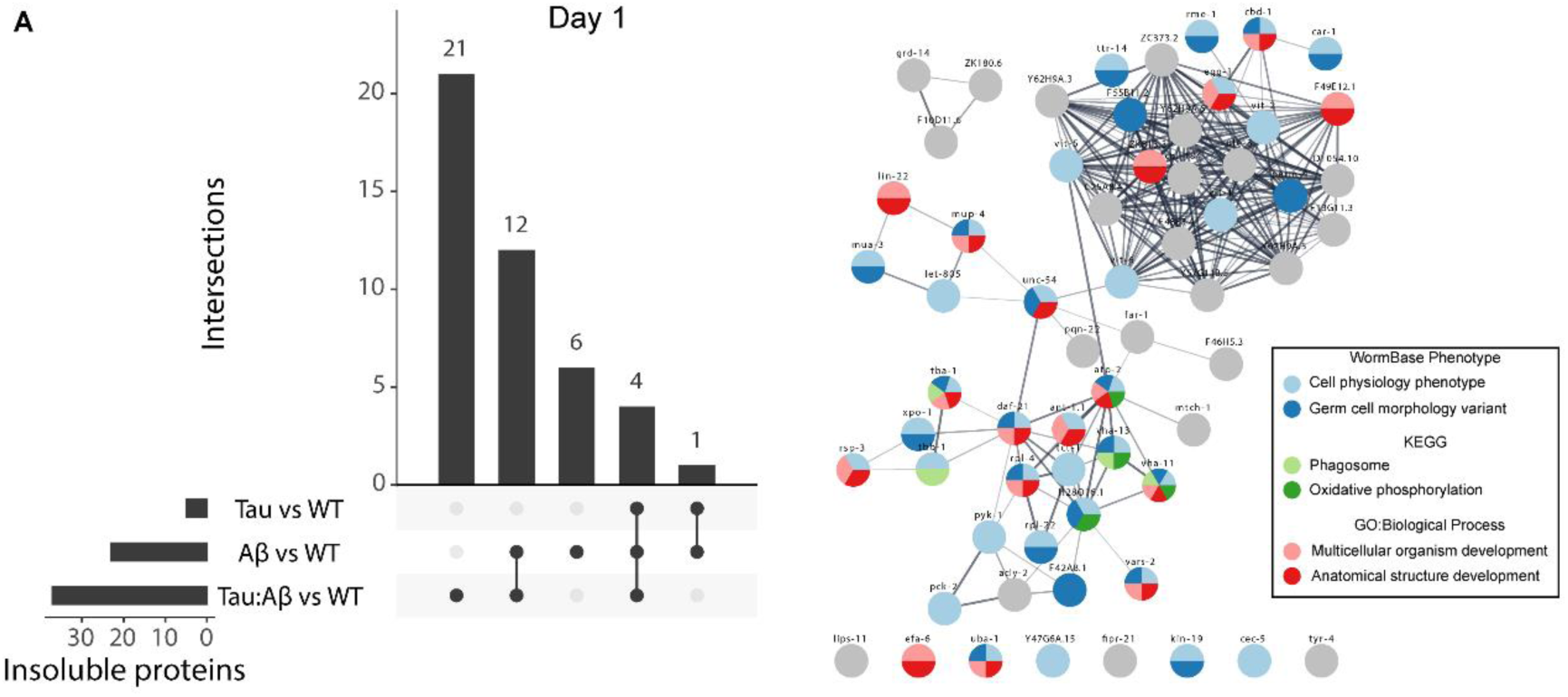
Expression of Tau and Aβ increase insoluble proteins early in life. (A) Upset plot of the number of insoluble proteins significantly increased in the AD strains compared to WT on day 1 of adulthood. The intersections are the number of proteins shared between the different comparisons, which are represented by a line connecting dots. q ≤ 0.05. (B) GO analysis of insoluble proteins increased in Tau;Aβ strain found significant increase of lipid transport genes, heatmap shows the abundance of those lipid transport genes (C) Protein-protein interaction network of insoluble proteins increased in the AD mutant strains compared to WT. 3 biological replicates, q ≤ 0.1.

To gain insights into the functional interactions of the insoluble proteins in the AD strains, we constructed a protein-protein interaction network. This network revealed associations with germ cells and cell physiology, oxidative phosphorylation, and organism development (Fig. 4A).

Notably, the expression of either Tau or Aβ alone led to increased insolubility of proteins associated with germ cell morphology and oxidative phosphorylation with potential relevance to the observed fertility defects and mitochondrial functional changes. Perturbation in mitochondrial function is also observed in human AD, suggesting that even at an early stage of adulthood, the Tau;Aβ expressing strain exhibits specific aspects relevant to AD^23,24^.

### Increased protein insolubility induced by Tau and Aβ expression in middle-aged animals

Building upon our observations of increased protein insolubility in the Tau;Aβ expressing strain in young animals, we investigated this molecular phenotype during aging. We collected insoluble proteins from day 3, 5, and 8 adult worms of the WT, Tau, Aβ, and Tau;Aβ expressing strains, followed by mass spectrometric analysis (see Supplementary Table 1-4). Our findings revealed that comparing insoluble proteins significantly elevated on day 3, 5, and 8 in the WT, Tau, Aβ, and Tau;Aβ strains compared to day 1 adults, we observed overlaps on day 3, 5 and 8 adults with day 1 adults for all four strains (Fig. 5A). The abundance of insoluble proteins in the WT strain increased with age (Fig. 5A), consistent with previous reports^18,25,26^. On day 5, the Tau expressing strain showed an increase in insoluble protein abundance compared to day 1. On day 8, the Aβ and Tau;Aβ strains exhibited an increase in insoluble proteins compared day 1. In contrast, the Aβ and Tau;Aβ strains exhibited a lower amount of insoluble proteins on day 3 and 5 (Fig. 5A).

**Figure 5:**
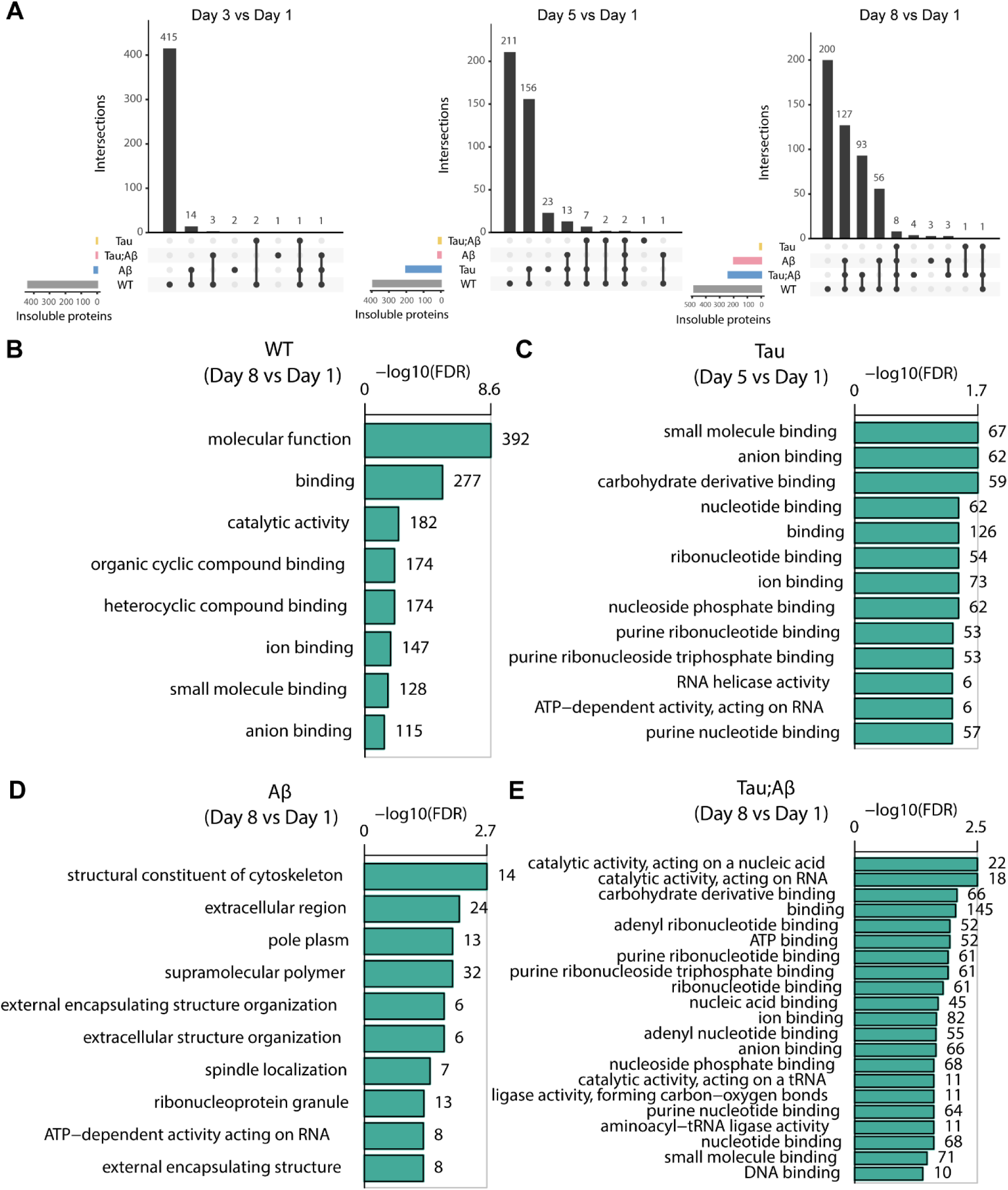
Expression of Tau and Aβ increases day 8 insoluble proteome. (A) Upset plot of the number of insoluble proteins significantly increased in the AD strains compared to WT on day 3 vs day 1, day 5 vs day 1, and day 8 vs day 1 of adulthood and the number of proteins shared between the different comparisons on each day. q ≤ 0.05. (B) g:Profiler GO molecular function of proteins with increased insolubility on day 8 vs day 1 in WT. (C) g:Profiler GO molecular function of proteins with increased insolubility on day 5 vs day 1 Tau. (D) Wormbase Gene ontology enrichment of proteins with increased insolubility on day 8 vs day 1 in Aβ. (E) g:Profiler GO molecular function of proteins with increased insolubility on day 8 vs day 1 in Tau;Aβ.

Enrichment analysis revealed an enrichment of proteins involved in ion and small molecule binding in the WT strain on day 8 compared to day 1 (Fig. 5B). Similarly, enrichment analysis of the Tau strain on day 5 compared to day 1 highlighted an increase in proteins associated with ribonucleotide binding (Fig. 5C). Enrichment analysis of the Aβ strain on day 8 compared to day 1 revealed an increase in proteins associated with cytoskeleton structure and ribonucleoprotein granules (Fig. 5D).

Furthermore, enrichment analysis of the Tau;Aβ expressing strain on day 8 compared to day 1 showed an increase in proteins involved in ribonucleotide binding (Fig. 5E). The increase in insolubility of ribonucleotide binding proteins may connect back to our previously mentioned observation of decreased nucleotide metabolites in our day 1 metabolomics data.

Supplementary Figure 4A demonstrates that the insoluble proteome of the Tau;Aβ strain was increased compared to the WT strain on day 3, 5, and 8, in comparison to Tau or Aβ alone. Enrichment analysis of the insoluble proteome on day 3 and day 5 of the Tau;Aβ strain compared to WT highlighted numerous ribosomal proteins (Supplementary Fig. 4B). When comparing day 3, 5, and 8 of the Tau;Aβ strain to day 1, the insoluble proteins observed on day 3 and 5 were also present on day 8 (Supplementary Fig. 4C). There was an overlap of insoluble proteins identified on day 1, 3, 5, and 8 of the Tau;Aβ strain compared to WT, with each day also featuring proteins not observed on the other days (Supplementary Fig. 4D).

The aging insoluble proteome analysis revealed that the Tau;Aβ expressing strain exhibited an early increase in insoluble proteins on day 1 (Fig. 4), which decreased after day 1 to then increase again on day 8. The aging insoluble proteome of the WT, Tau, and Tau;Aβ strains consisted of proteins involved in binding a range of small molecules.

Taken in total, our data suggests that the expression of Tau and/or Aβ leads to a complex remodeling of protein-wide insolubility. The non-linear aging response observed may reflect compensatory mechanisms aimed at preventing or removing insoluble proteins.

### Tau and Aβ expression in young adult worms resemble WT aging at insoluble proteome and metabolic levels

Comparing insoluble proteome and metabolomic changes in the day 1 Tau;Aβ expressing strain to the aged WT strain revealed similarities between the two conditions. Proteins that were more abundant in the insoluble proteome of day 8 WT compared to day 1 WT overlapped with the majority of insoluble proteins in day 1 Tau;Aβ compared to day 1 WT (Fig. 6A). These overlapping proteins are involved in lipid transport and ribosome function. Similarly, some of the metabolites that were altered in day 8 WT compared to day 1 WT also overlapped with metabolites altered in day 1 Tau;Aβ compared to day 1 WT (Fig.6B). These overlapping metabolites fall mostly in the categories of lipids and amino acids (Fig.6C). In the Tau;Aβ strain, these metabolites had elevated abundance levels on day 1, resembling the levels observed in day 8 WT (Fig. 6C).

**Figure 6:**
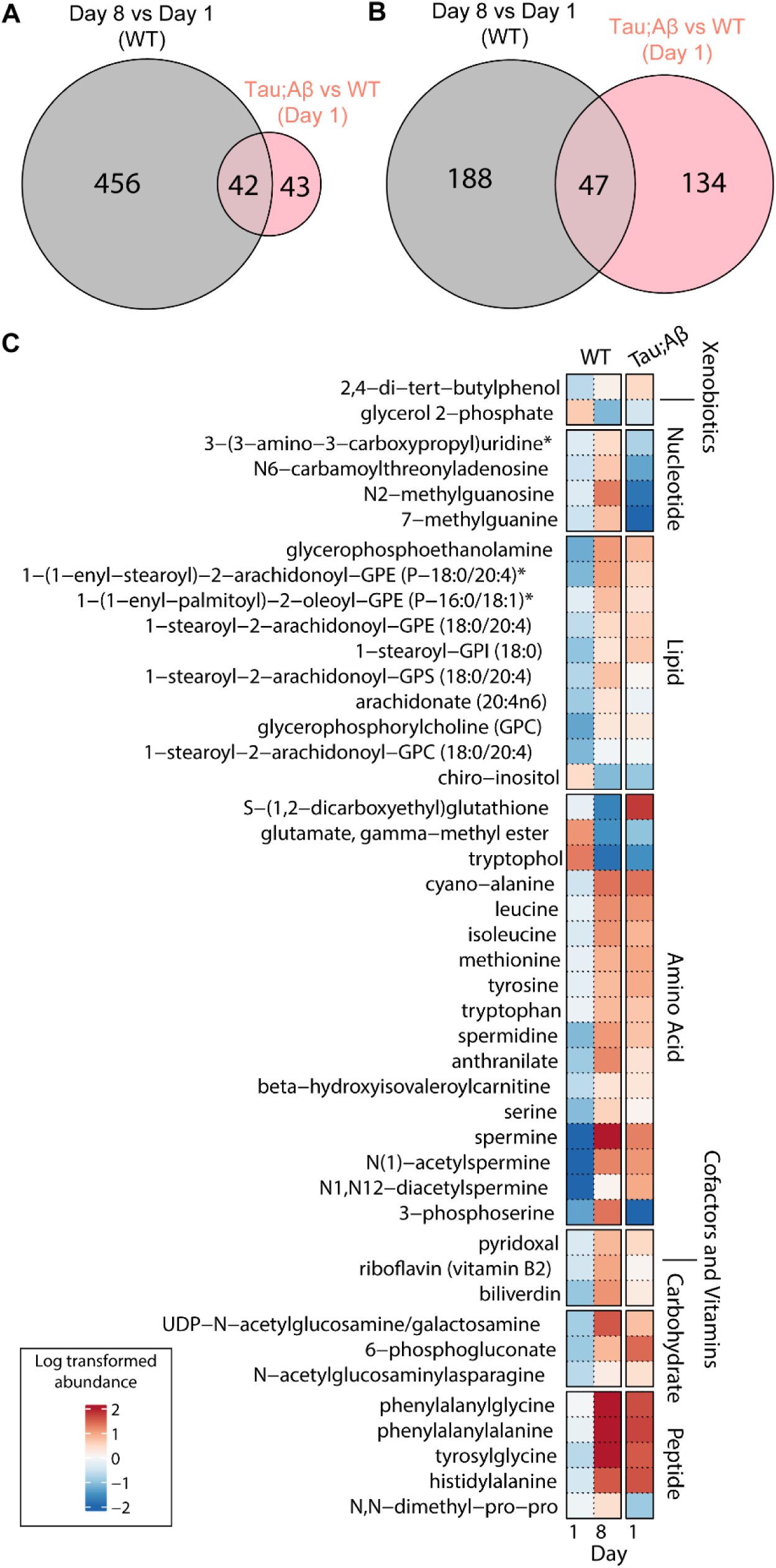
Tau and Aβ expression in young adult worms resembles WT aging at insoluble proteome and metabolomic levels. (A) Venn diagram of insoluble proteins that are increased on day 1 Tau;Aβ vs day 1 WT compared to day 8 WT vs day 1 WT and GO biological process of the insoluble proteins shared between the day 8 WT and day 1 Tau;Aβ. (B) Venn diagram of the metabolites in day 1 Tau;Aβ vs day 1 WT compared to day 8 WT vs day 1 WT and (C) a heatmap of the shared metabolites.

This suggests that the expression of Tau and Aβ in young adult worms resembles the aging-associated changes observed in the insoluble proteome and metabolic pathways of WT worms. The simultaneous expression of Tau and Aβ in young adult worms appears to accelerate the aging process, particularly those affecting lipid pathways. The observed similarities between the insoluble proteome and metabolic changes in the Tau;Aβ strain on day 1 and the aged WT strain suggest that the presence of Tau and Aβ proteins leads to accelerated aging on a proteomic and metabolomic level, resembling the changes typically seen in older organisms. These alterations in lipid pathways may contribute to the dysregulation of cellular processes and potentially exacerbate the detrimental effects associated with aging.

## Discussion

We developed a *C. elegans* model of Alzheimer’s disease and Alzheimer’s related disease (ADRD) proteotoxicity by expressing both human Aβ and human 4R1N Tau containing the V337 mutation in neurons. The generation of this new dual strain facilitated an investigation into the combined effects of Aβ and Tau expression on disease progression through phenotypic and geromic analysis.

The simultaneous expression of Tau and Aβ has been demonstrated to influence various phenotypes encompassing progeny viability, body movement, lifespan, and neurol loss^11,12^. In our Tau;Aβ strain, we observed a synergistic effect on health (body movement and lifespan). Intriguingly, mitochondrial function and fertility phenotypes were predominantly influenced by either Tau or Aβ alone. This dichotomy suggests that the combined expression of Tau and Aβ may have exert distinct effects on different phenotypes.

Collectively, these findings provide compelling evidence for the intricate relationship between Aβ and Tau expression and their associated phenotypic outcomes, underscoring the significance of their interplay in the context of neurodegenerative diseases.

The phenotypic findings give rise to further questions, specifically whether Tau and Aβ drive different aspects of the disease in humans or if all aspects of the disease result from a combined effect. Additionally, there is a pressing need to delve deeper to elucidate the mechanisms involved and navigating whether Tau and Aβ are directly interacting. These avenues of inquiry are pivotal for comprehensive understanding of the molecular intricacies underpinning ADRD pathogenesis.

To delve further into the intricate effects of Tau and Aβ we conducted a geromic analysis, including transcriptomics, metabolomics, and insoluble proteomics. In addition, we were interested in determining the temporal order of the phenotypic versus the multi-omic changes. Transcriptomic analysis of the young adult Tau;Aβ strain highlighted significant changes in mRNAs related to reproduction and mitochondrial function, potentially linking to our observed phenotypes (Supplementary Fig. 5). Notably, we identified similarities with the transcriptomic data published by Wang et al., showing shared upregulated mRNAs and genes associated with protein phosphorylation and UDP-glucuronosyl transferase^11^. However, differences were noted in downregulated genes possibly attributed to the variation in the Tau strain used in creating the Tau;Aβ strain. This underscores the importance of employing multiple models with different protein isoforms and mutations to comprehensively understand the range of transcriptomic changes in the disease.

Metabolomics analysis revealed early-life increases in metabolite levels for the Tau;Aβ strain, particularly lipids and amino acids. Elevated amino acid levels can be attributed to higher energy demand, prompting amino acid oxidation as an alternative energy source^27^. Furthermore, elevated amino acid levels may be connected to increased protein synthesis and activation of mTOR inhibiting autophagy^27^. This underscores the importance of studying a range of omics, as a synergistic effect between Tau and Aβ was specifically observed at the metabolomic level, emphasizing need to explore various omics to avoid potential oversights.

The insoluble proteome analysis indicated an increase in insoluble proteins in the young adult Tau;Aβ strain compared to the WT strain, associated with germ cell morphology and oxidative phosphorylation, which may also impact our observed phenotypes. These changes align with the increase in amino acids observed in our metabolomic data and its connection to protein synthesis. Additionally, alterations in mRNAs related to ribonucleotide binding and ribosome structure which could associated with the increase in insoluble proteins.

Thus far we have observed that phenotypic changes and multi-omic changes both occur in early life with simultaneous expression of Tau and Aβ. Given these early life changes we observed we were keen on examining whether these alterations persist as the organisms age. Our observations revealed a progressive increase in metabolite alterations associated with energy and lipid metabolism in the Tau;Aβ strain, mirroring changes observed in AD^23,24,28–31^. These findings imply that the simultaneous expression of Tau and Aβ induces disruptions in energy metabolism, lipid metabolism, and mitochondrial function, manifesting early in life and that persists with age. Significantly, this finding prompts consideration of whether metabolomic dysfunction represents one of the initial indicators of the disease.

Additionally, aging in the Tau;Aβ strain was accompanied by an increase in insoluble proteins associated with ribonucleotide and small molecule binding. Dysfunction in these binding proteins may prevent proper binding, leading to their aggregation with age, a phenomenon observed in the initial stages of AD^32^. Notably, in the young adults Aβ appears to drive protein insolubility, yet this trend does not persist with age. The observed plateau and subsequent increase in insolubility at day 8 could be attributed to the induction of Aβ at the late larval stage, resulting in increased protein insolubility at day 1. This may represent an acute stress response that does not have a prolonged effect beyond the first 24 hours of expression. However, this initial stress may have lasting effects, becoming evident in the increased insoluble proteome at the advanced age of day 8.

Comparative analysis of the insoluble proteome and metabolomic changes between the day 1 Tau;Aβ strain and the aged WT strain unveiled striking similarities, suggesting a convergence of molecular alterations. The overlapping insoluble proteins predominantly associated with lipid transport and ribosome function, along with overlapping metabolites already elevated in the day 1 Tau;Aβ strain, became more abundant with age in the WT strain. The simultaneous expression of Tau and Aβ in young adult worms appears to accelerate and intensify aging processes, particularly those impacting lipid pathways. These disruptions in lipid pathways may contribute to the dysregulation of cellular processes, potentially exacerbating the detrimental effects associated with aging.

The Tau;Aβ strain, when juxtaposed with the findings from the prior studies by Wang et al. and Benbow et al., collectively illustrates that the concurrent expression of both Aβ and Tau gives rise to distinct neuronal and systemic phenotypes^11,12^. This integrated model not only advances our understanding of the individual contributions of Tau and Aβ but also illuminates their synergistic effects, providing a comprehensive perspective on their impact on AD progression from early life. It facilitates the examination of early systemic changes in AD, and aids in the identification of potential therapeutic targets.

Crucially, the Tau;Aβ strain contributes valuable insights into AD by unveiling transcriptomic changes related to reproduction and mitochondrial function, significant shifts in metabolomics, and alterations in the insoluble proteome. These findings not only align with, but also expand upon previous studies, underscoring the intricate relationship between Aβ and Tau expression and their consequential impact on neurodegenerative processes. Furthermore, our research accentuates the convergence of molecular alterations between the Tau;Aβ strain and aging in the WT strain, particularly within lipid pathways.

In summary, the Tau;Aβ strain, in conjunction with earlier studies, provides a holistic portrayal of early systemic changes in Alzheimer’s disease. Our study introduces additional transcriptomic timepoints, complemented by metabolomic and insoluble proteomic data, contributing to the existing field of Tau and Aβ co-expression *C. elegans* models. Moreover, our results validate prior studies that observed AD-associated proteins affected reproduction, lifespan, and body movement in a *C. elegans* model expressing both Tau and Aβ^11,12^. Our study yields valuable insights into potential therapeutic targets and serves as a resource model for studying AD pathogenesis. By uncovering changes at the metabolomic and insoluble proteomic levels, our research introduces novel perspectives to the field, providing geromic datasets that can be mined to enhance our understanding of ADRD progression.

## Methods

### *C. elegans* strains

The transgenic strains utilized in this study include non-transgenic wild-type N2, CK10 (P*_aex-3_::*h4R1N Tau V337M; P*_my_*_o_*_-2_*::GFP)^14^, CL2355(*smg-1*(*cc546);*P*_snb-1_*::Aβ_1-42_ + P*_mtl-2_*::GFP)^13^, and the simultaneous expression strain GL405 (P*_aex-3_*::h4R1N Tau V337M; P*_myo-2_::*GFP; *smg-1*(*cc546)*;P*_snb-1_*::Aβ_1-42_ + P*_mtl-2_*::GFP). CL2355 was obtained from the *Caenorhabditis* Genetic Center and CK10 was obtained from the lab of Dr. Brian Kraemer at the University of Washington. N2, CK10, CL2355, GL405 strains are maintained at 15°C due to the *smg-1*(*cc546)* gene, which allows for the temperature-dependent expression of Aβ. Once the worms reached the fourth larval stage (L4), they were shifted to 25°C to induce Aβ expression. Throughout the study, the worms were cultured on nematode growth media (NGM) (containing: Becton Dickinson and Company Difco Agar, Bacteriological, ref:214510, Gibco Bacto Peptone Enzymatic Digest of Protein ref: 211820, Sigma-Aldrich Sodium Chloride S9888-1KG, Sigma-Aldrich Magnesium Sulfate M7506-500G lot#SLBR0877V, Sigma-Aldrich Calcium Chloride C1016-500G lot#SLCC1966) plates (VWR, petri dish polystyrene disposable sterilized size: 60 x 15mm, cat no.25384-090) containing *E. coli* OP50 and passaged once a week.

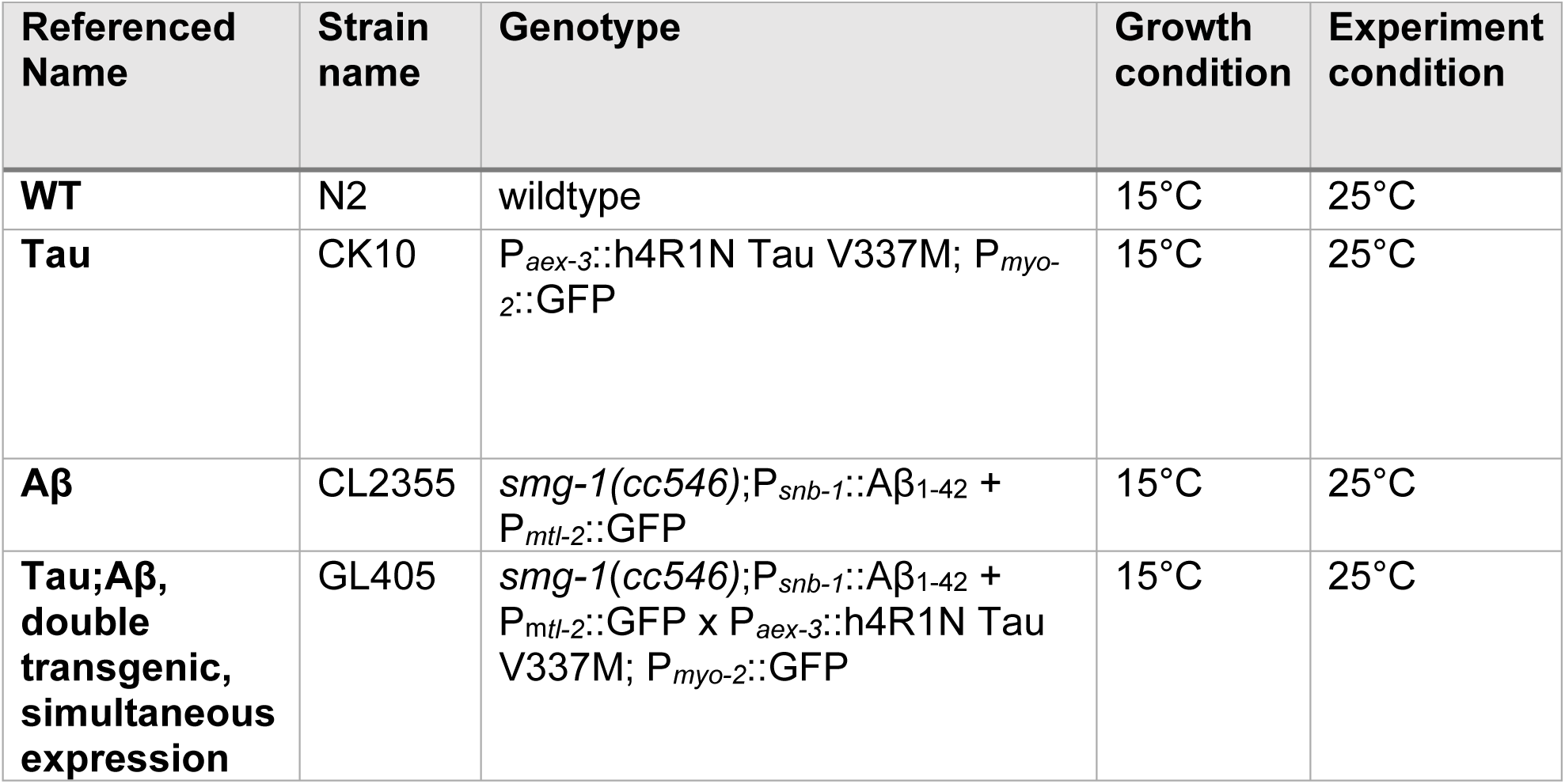

### Hermaphrodite self-fertility and fecundity assay

Egg lays were conducted at 15°C for each strain. Once worms develop to the L4 stage, 10 hermaphrodite worms were moved to individual small NGM plates (Gen Clone, tissue culture dishes 38x13mm, cat# 25-200) and shifted to 25°C. On day 1 of the adult stage, the number of eggs laid was counted to determine fecundity and the number of hatched larvae was counted to determine fertility. The worms were then transferred to fresh NGM plates. The number of eggs laid, and hatched larvae were counted each day until the end of their reproductive cycle. To determine if there were significant differences in fertility and fecundity among the strains, statistical analysis was performed using ordinary one-way ANOVA with Tukey’s multiple comparisons test and the data consisted of 3 biological replicates.

### Thrashing assay

Following egg lays conducted at 15°C, the worms were allowed to develop to the L4 stage and then transferred to 10 µg/mL FUdR plates, followed by a shift to 25°C. On day 1 of the adult stage, a droplet of S-basal solution (containing: Sigma-Aldrich Potassium Phosphate Monobasic P5379-1KG lot#SLCG1814, Sigma-Aldrich Potassium Phosphate Dibasic P3786-1KG lot#SLCC4706, and Sigma-Aldrich Sodium Chloride S9888-1KG) was placed on a glass slide, and a single worm was placed into the droplet to acclimate for 30 seconds. Subsequently, the body bends of the worm were counted for another 30 seconds. This process was repeated for each strain, with 10-20 worms included in each of the 3 biological replicate. To determine if there were significant differences in the number of body bends between the strains, statistical analysis was performed using ordinary one-way ANOVA with Tukey’s multiple comparisons test and the data consisted of 3 biological replicates.

### Mitochondrial oxygen consumption rate assay

To evaluate mitochondrial function, a mitochondrial oxygen consumption rate (OCR) assay was performed. Over 200 worms were mass cultured on 100 mm NGM agar plates (petri dish polystyrene disposable sterilized size: 100 x 15mm, cat no. 25384-342) at 15°C. Age synchronization was achieved by isolating eggs through hypochlorite treatment of gravid adults. Once the worms reached the L4 stage, they were transferred to NGM plates containing 10 µg/mL FUdR and shifted to a temperature of 25°C. Approximately 200 worms were collected on day 1 of the adult stage for further analysis.

An Agilent Seahorse XFe96 Analyzer was utilized to measure the OCR using a previously described protocol^33^. Briefly, for each strain, approximately 20 hermaphrodite worms were loaded into each well of a 96-well microtiter plate (Agilent Seahorse XFe96/XF Pro FluxPak, ref#103792-100), with 8 technical replicates per strain. Basal OCR was measured for 5 cycles, where each cycle consisted of 2 minutes of mixing in the well, 4 minutes of waiting, and 2 minutes of OCR measurement in the well.

To determine if there were significant differences in OCR between the strains, statistical analysis was performed using ordinary one-way ANOVA, followed with Tukey’s multiple comparisons test and the data consisted of 3 biological replicates.

### Lifespan assay

To assess the lifespan of the worms, egg lays were conducted at 15°C. Once the worms reached the L4 stage, approximately 40-60 worms per strain were transferred to NGM plates containing 10 µg/mL FUdR and maintained at a temperature of 25°C. Three technical replicates were performed for each strain to ensure robustness and reproducibility of the results. The worms were monitored every two days for touched provoked movement. This was achieved by gently poking the worms with a platinum wire and observing their response. The absence of movement upon stimulation was considered an endpoint, indicating mortality.

Statistical analysis was conducted using survival representation (Kaplan-Meier) in GraphPad Prism™ software. Survival curves were generated, and the lifespan of each strain was compared using the Log-rank (Mantel-Cox) test. This analysis allowed for the assessment of significant differences in survival between strains and the data consisted of 3 biological replicates.

## 3’ Tag RNA-seq

For the 3’ Tag RNA-seq analysis, worms were mass cultured at 15°C, and age-synchronization was achieved by isolating eggs through hypochlorite treatment of gravid adults. Once the worms reached the L4 stage, they were transferred to FUdR NGM plates and maintained at a temperature of 25°C.

Approximately 1000 worms and 3 biological replicates per strain were collected on the first day of the adult stage and stored in a 1.5 mL Eppendorf tube containing 300-500 µL of RNA lysis buffer from the Zymo Research RNA extraction kit (cat#R1055). The samples were immediately placed at -80°C for storage. To facilitate RNA extraction, the samples underwent three cycles of freeze-thaw, alternating between room temperature and -80°C. Subsequently, the samples were incubated on a heat block at 55°C for 0.5-1 hour.

RNA extraction was performed using a Zymo Research RNA extraction kit according to the provided protocol. The protocol included a DNAse treatment step to remove any genomic DNA contamination. The elution step resulted in approximately 40 µL of purified RNA.

The prepared RNA samples were then sent out on dry ice for 3’ Tag RNA-Seq analysis. The sequencing was conducted at the DNA Technologies and Expression Analysis Cores at the UC Davis Genome Center, with support from NIH Shared Instrumentation Grant 1S10OD010786-01.

### Binarized transcriptomic aging (BiT age) clock calculation

Read counts from the RNA-seq data were transformed into counts per million using edgeR ^34^. Using this file and the regression coefficients in https://github.com/Meyer-DH/AgingClock/blob/main/Data/Predictor_Genes.csv we calculated the biological age of each sample from day 1 using the Python script https://github.com/Meyer-DH/AgingClock/blob/main/src/biological_age_prediction.py

### Metabolomics

For metabolomic analysis, worms were mass cultured at 15°C and synchronized by isolating eggs using hypochlorite treatment. Upon reaching the L4 stage, the worms were transferred to FUdR 100mm NGM plates at 25°C. At the adult stage day 1,3,5, and 8, approximately 250µl of worms and 3 biological replicates were collected by washing off with S-basal solution and immediately snap freezing ^35,36^. These samples were then submitted to Metabolon Inc. (Morrisville, NC, USA) for global metabolic profiling.

Sample preparation was conducted using the automated MicroLab STAR® system from the Hamilton Company. Prior to the extraction process, several recovery standards were added to the samples for quality control purposes. The extracted samples were analyzed using a Waters ACQUITY ultra-performance liquid chromatography (UPLC) system and a Thermo Scientific Q-Exactive high resolution/accurate mass spectrometer equipped with a heated electrospray ionization (HESI-II) source and Orbitrap mass analyzer operating at a mass resolution of 35,000.

Raw data obtained from the analysis was processed by Metabolon’s hardware and software. This involved extraction of the data, identification of peaks, and quality control processing. Each metabolite was corrected by registering the medians to a value of one (1.00) and normalizing each data point proportionately. In certain cases, metabolite data was further normalized to total protein levels determined by Bradford assay to account for differences in metabolite levels due to variations in the amount of biological material present in each sample.

A total of 767 biochemical features were identified, including 695 metabolites with known composition and 72 compounds with unknown structural identity. Statistical analysis was performed using two-way ANOVA to identify biochemicals that exhibited significant differences between experimental groups. Additionally, a false discovery rate (q-value) was estimated to account for multiple comparisons.

### Insoluble proteome

For the insoluble proteome analysis, which is based on ^25^, worms were mass cultured at 15°C and age-synchronized by isolating eggs using hypochlorite treatment of gravid adults. Upon reaching the L4 stage, the worms were transferred to FUdR NGM plates at 25°C. Approximately 3,000 worms at the adult stage day 1,3,5 and 8 were collected in S-basal solution and stored at -80°C.

To extract the insoluble protein fraction, the samples were thawed and vortexed with worm lysis buffer containing 20 mM Tris base (pH 7.4), 100 mM NaCl, 1 mM MgCl_2_, and EDTA-free protease inhibitor. The lysate was then sonicated for 10 cycles and centrifuged at 3,000 x g for 4 minutes in a cold room. The resulting supernatant, containing the aqueous-soluble protein fraction, was transferred to new 1.5 mL Eppendorf tubes, and the protein concentration was quantified using a BCA assay.

The protein lysate was further processed by centrifuging it for 15 minutes at 20,000 x g in 4⁰C. The resulting supernatant was saved as the SDS-soluble fraction. The pellet obtained from this step was washed with 500µl of worm lysis buffer containing 1% SDS and centrifuged at 20,000 x g for 15 minutes. The supernatant was removed, and this washing step was repeated two more times to remove the SDS-soluble fraction. The remaining pellet, known as the 1% SDS-insoluble protein fraction, was resuspended in 60µl of 7% formic acid (Thermo Scientific, ref: 85178) and vortexed to dissolve the pellet. Subsequently, the sample was sonicated for 30 minutes and dried in a vacuum concentrator for 1 hour to remove the formic acid solution.

The total amount of 1% SDS-insoluble protein, per sample, was prepared for trypsin digestion. Each sample was reduced using 20 mM dithiothreitol in 50 mM triethylammonium bicarbonate buffer (TEAB) at 50°C for 10 min, cooled to room temperature (RT) and held at RT for 10 min, and alkylated using 40 mM iodoacetamide in 50 mM TEAB at RT in the dark for 30 min. Samples were acidified with 12% phosphoric acid to obtain a final concentration of 1.2% phosphoric acid. S-Trap buffer consisting of 90% methanol in 100 mM TEAB at pH ∼7.1, was added and samples were loaded onto the S-Trap mini spin columns. The entire sample volume was spun through the S-Trap mini spin columns at 4,000 × g and RT, binding the proteins to the mini spin columns. Subsequently, S-Trap mini spin columns were washed twice with S-Trap buffer at 4,000 × g at RT and placed into clean elution tubes. Samples were incubated for one hour at 47 oC with sequencing grade trypsin (Promega, San Luis Obispo, CA) dissolved in 50 mM TEAB at a 1:25 (w/w) enzyme:protein ratio. Afterwards, trypsin solution was added again at the same ratio, and proteins were digested overnight at 37°C.

Peptides were sequentially eluted from mini S-Trap spin columns with 50 mM TEAB, 0.5% formic acid (FA) in water, and 50% acetonitrile (ACN) in 0.5% FA. After centrifugal evaporation, samples were resuspended in 0.2% FA in water and desalted with C18 Ziptips (MilliporeSigma, Burlington, MA). Desalted peptides were then subjected to an additional round of centrifugal evaporation and re-suspended in 20 µL of 0.2% FA in water and 1 µL of indexed Retention Time Standard (iRT, Biognosys, Schlieren, Switzerland). Samples were then subjected to mass spectrometric analysis using a high-performance liquid chromatography (HPLC) system combined with a chip-based HPLC system (Eksigent nano-LC) directly connected to a quadrupole time-of-flight mass spectrometer (TripleTOF 5600, a QqTOF instrument) as detailed in a step-by-step protocol by Xie et al ^25^.

Each sample was acquired in data-dependent acquisition (DDA) mode to build peptide spectral libraries, as described in Xie et al ^25^. Data-Independent Acquisition (DIA)/SWATH data was processed in Spectronaut (version 14.10.201222.47784) using DIA. Data extraction parameters were set as dynamic and non-linear iRT calibration with precision iRT was selected. DIA data was matched against an in-house *Caenorhabditis elegans* spectral library that provides quantitative DIA assays for 3,651 *C. elegans* peptides corresponding to 910 protein groups and supplemented with scrambled decoys (library size fraction of 0.01), using dynamic mass tolerances and dynamic extraction windows. DIA/SWATH data was processed for relative quantification comparing peptide peak areas from different days. Identification was performed using 1% precursor and protein q-value. Quantification was based on the peak areas of extracted ion chromatograms (XICs) of 3 – 6 MS2 fragment ions, specifically b- and y-ions, with q-value sparse data filtering and iRT profiling being applied (Supplementary Table 1-4). For this sample set, local normalization was not implemented. Differential protein expression analyses for all comparisons were performed using a paired t-test, and p-values were corrected for multiple testing, using the Storey method ^37^. Specifically, group wise testing corrections were applied to obtain q-values. Protein groups with at least two unique peptides, q-value ≤ 0.01, and absolute Log2(fold-change) > 0.58 are significantly altered.

### Protein-protein interaction network

We extracted 62 proteins with increased abundance (q-value < 0.1) in the insoluble fraction compared to N2 controls at day 1. We loaded this list of proteins into Cytoscape ^38^ and generated a protein-protein interaction network using the stringApp plugin ^39^.

Then, we performed enrichment analysis for Biological Processes (Gene ontology), KEGG pathways, and WormBase phenotypes. We colored the nodes of proteins from the top-enriched processes in the network.

### Geromic analysis

We conducted several omic analyses to gain a comprehensive geromic understanding of the molecular changes associated with the studied conditions. The following approaches were employed:

#### Enrichment analysis

For the RNAseq and insoluble proteome data, we performed enrichment analysis using g:Profiler. The custom background used for each specific comparison included all genes or all proteins measured in that particular analysis. The enrichment analysis was carried out with a user threshold of 0.05 and significance was determined using the Benjamini-Hochberg false discovery rate (FDR) correction. Additionally, WormBase Enrichment Analysis was utilized for some of the RNAseq enrichment, applying a q-value threshold of 0.05 and the previously mentioned custom background. Metaboanalyst was utilized for enrichment analysis of metabolomics data, employing a custom background consisting of all measured metabolites.

#### WGCNA (weighted gene co-expression network analysis)

To identify modules of metabolites that exhibited correlated patterns across samples, we employed WGCNA. This analysis involved using the abundance data of metabolites and allowed us to identify groups or modules of metabolites that displayed similar expression profiles, providing insights into potential coordinated metabolic pathways or regulatory mechanisms^40^.

#### Generation of heatmap and dot plots

RStudio was utilized to generate heatmaps and dot plots to provide a comprehensive visual overview of observed molecular changes and enriched pathways. Code and referenced files can be found at https://github.com/Holcom-AM/multi-omic-analysis and https://data.mendeley.com/datasets/zc4s3dszf5/1,

## Resource Availability

### Lead contact

Further information and requests for resources and reagents should be directed to and will be fulfilled by the lead contact, Gordon J. Lithgow.

### Materials availability

The Tau;Aβ strain generated in this study is available from the lead contact upon request.

### Data and code availability

Raw data and complete MS data sets have been uploaded to the Mass Spectrometry Interactive Virtual Environment (MassIVE) repository, developed by the Center for Computational Mass Spectrometry at the University of California San Diego, and can be downloaded using the following link: https://massive.ucsd.edu/ProteoSAFe/private-dataset.jsp?task=e2c3bbad2f214e88bc70ea487d71a3ed (MassIVE ID number: MSV000092031; ProteomeXchange ID: PXD042484. Enter the username and password in the upper right corner of the page: Username: MSV000092031_reviewer; Password: winter. Transcriptomics and metabolomics data can be found at https://github.com/Holcom-AM/multi-omic-analysis and https://data.mendeley.com/datasets/zc4s3dszf5/1.

## Supporting information

Supplementary Table 1 insoluble proteome

## Acknowledgements

We would like to thank all members of the Lithgow, Andersen, Schilling, and Furman labs. This work was supported by the Larry L. Hillblom Foundation, NIH Shared Instrumentation Grant 1S10OD010786-01, 1S10OD016281-01 (to Buck Institute for the TripleTOF mass spectrometry system), NIA grants RF1AG057358-01 and R01AG029631-10. This work was also supported by U54 AG075932 (PI Campisi/ MPI Schilling) and P01AI153559 (to Buck Bioinformatics Core). Traineeship supported by NIA T32 AG052374 awarded to AH. The authors thank Dr. Akos Gerencser and former Brand lab members for assistance with the Seahorse XFe96. The Aβ strain CL2355 is from the lab of Dr. Chris Link and the Tau strain CK10 is from the lab of Dr. Brian Kraemer. DeepVenn was used for venn diagrams and Biorender was used for the Fig 1A.

## Author contributions

Conceptualization: R.S., J.K.A, G.J.L.; Phenotypic analysis: R.S., A.H.; Transcriptomics: A.H.; Metabolomics: A.H.; Insoluble proteome: R.S., C.D.K., H.O., A.F., D.B., B.S.; Data analysis: M.F., A.H., C.D.K.; Writing: A.H. wrote the manuscript, with input from all authors; Supervision: J.K.A., G.J.L., D.F., B.S. M.F is co-first author.

---

**Extended Data Figure 1:**
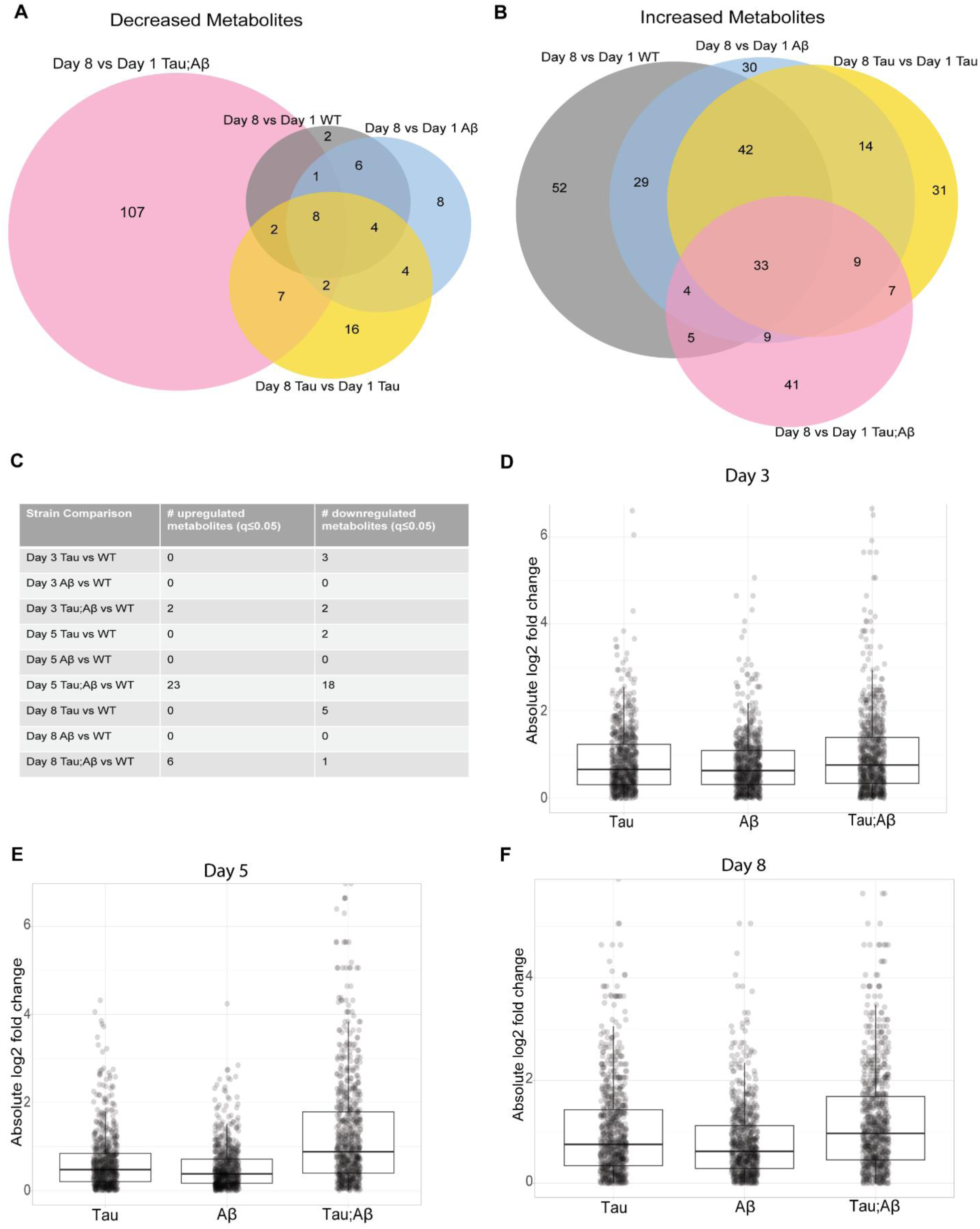
Enhanced Age-Related Metabolomic Changes in Tau;Aβ. (A) Venn diagram of downregulated metabolites on day 8 of adulthood in Tau, Aβ, and Tau;Aβ compared to day 1, significance was based on q ≤0.05. (B) Venn diagram of metabolites with increased abundance on day 8 of adulthood in Tau, Aβ, and Tau;Aβ compared to day 1, significance was based on q ≤0.05. (C) Table of number of significantly increased and decreased metabolites on day 3, 5 and 8 of adulthood in Tau, Aβ, and Tau;Aβ compared to WT, significance was based on q ≤0.05. (D) Bar graph of fold metabolite change in day 3 Tau, Aβ and Tau;Aβ compared to WT. (E) Bar graph of fold metabolite change in day 5 Tau, Aβ and Tau;Aβ compared to WT. (F) Bar graph of fold metabolite change in day 8 Tau, Aβ and Tau;Aβ compared to WT.

**Extended Data Figure 2:**
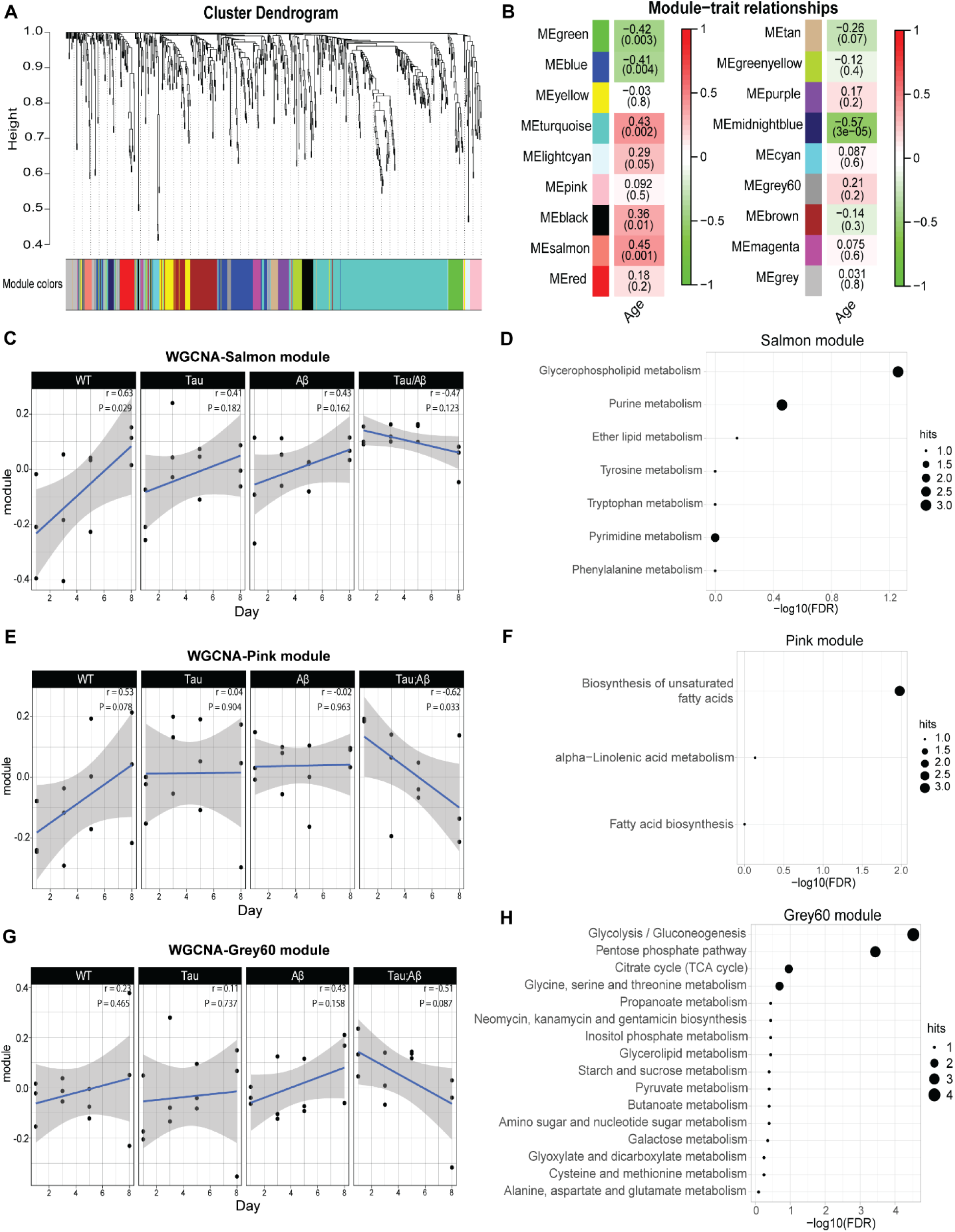
WGCNA of metabolomic changes. (A) Clustering dendrogram of metabolites with assigned module colors. (B) Module-trait relationship plot with rows representing the modules and column the trait of interest, which is age. Each cell contains the correlation and below is the p-value in parenthesis. The table’s cells are color-coded based on correlation, as indicated by the color legend. (C) WGCNA eigengene plot of salmon module with all 4 strains from day 1 to day 8. (D) Dot plot of MetaboAnalyst KEGG enrichment analysis of salmon module. (E) WGCNA eigengene plot of pink module with all 4 strains from day 1 to day 8. (F) Dot plot of MetaboAnalyst KEGG enrichment analysis of pink module. (G) WGCNA eigengene plot of grey60 module with all 4 strains from day 1 to day 8. (H) Dot plot of MetaboAnalyst KEGG enrichment analysis of grey60 module.

**Extended Data Figure 3:**
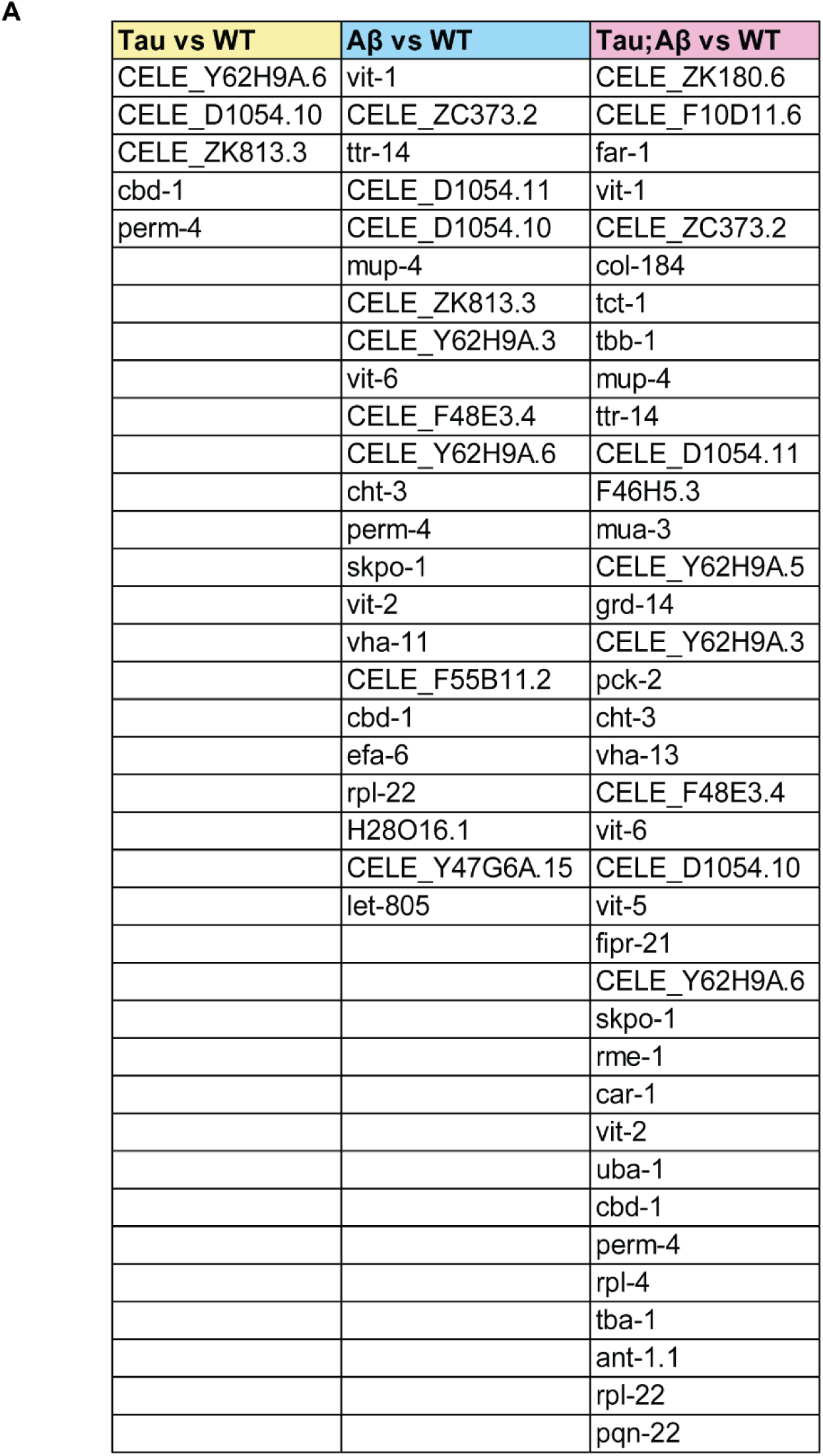
Increased abundance of insoluble proteome with Aβ expression. (A) List of insoluble proteins with increased abundance in Tau vs WT, Aβ vs WT and Tau;Aβ vs WT, significance was based on q ≤0.05.

**Extended Data Figure 4:**
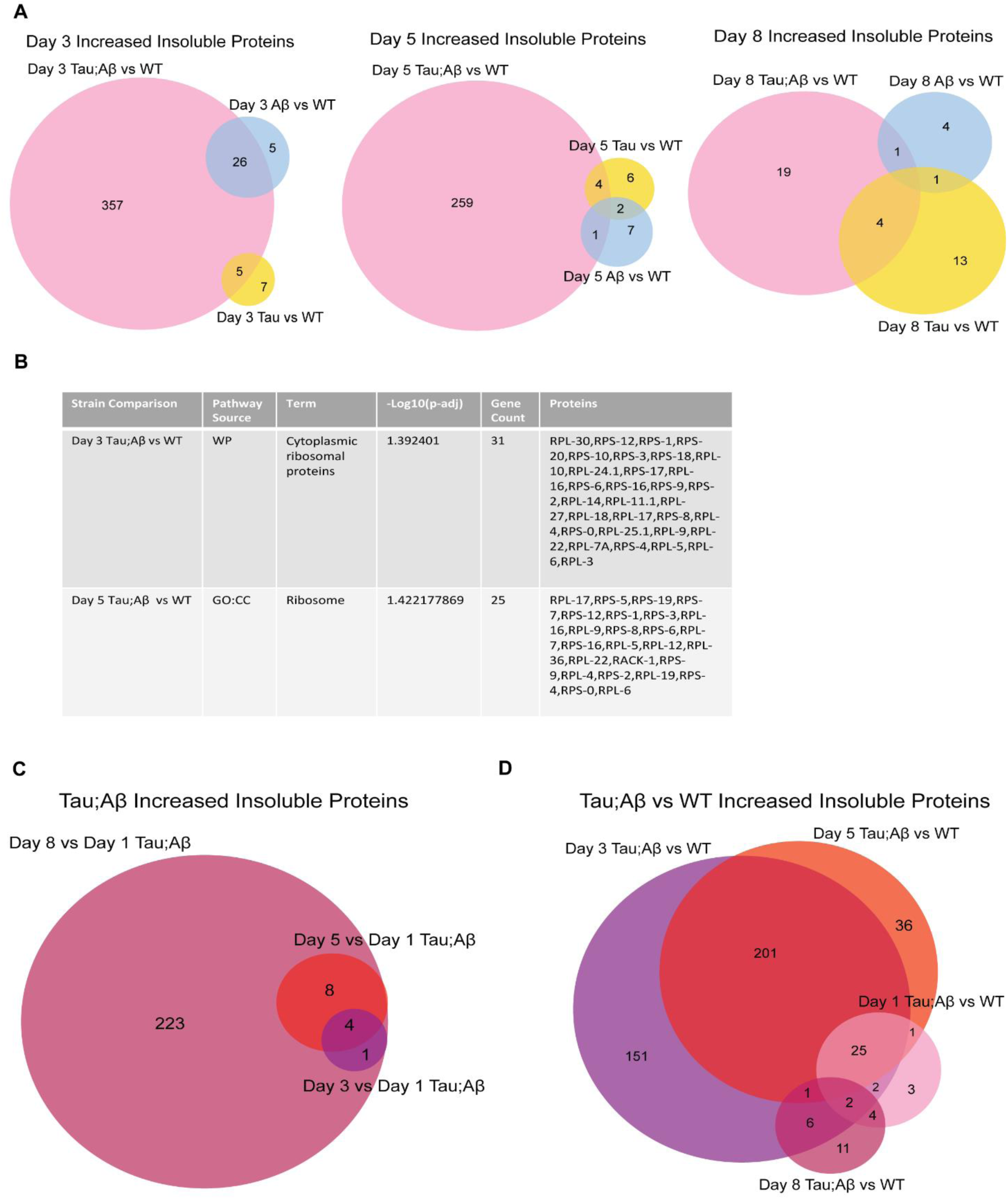
Alterations in aged insoluble proteome with enrichment for ribosomal proteins in Tau;Aβ strain. (A) Venn diagrams of insoluble proteins with increased abundance on day 3, 5 and 8 of adulthood in Tau, Aβ, and Tau;Aβ compared to WT, significance was based on q ≤0.05. (B) Table of g:Profiler enrichment of day 3 Tau;Aβ vs WT and day 5 Tau;Aβ vs WT. (C) Venn diagram of insoluble proteins that are increased on day 3, 5 and 8 of adulthood in Tau;Aβ compared to day 1 Tau;Aβ, significance was based on q ≤0.05. (D) Venn diagram of insoluble proteins increased on day 1, 3, 5 and 8 of adulthood in Tau;Aβ compared to WT, significance was based on q ≤0.05.

**Extended Data Figure 5:**
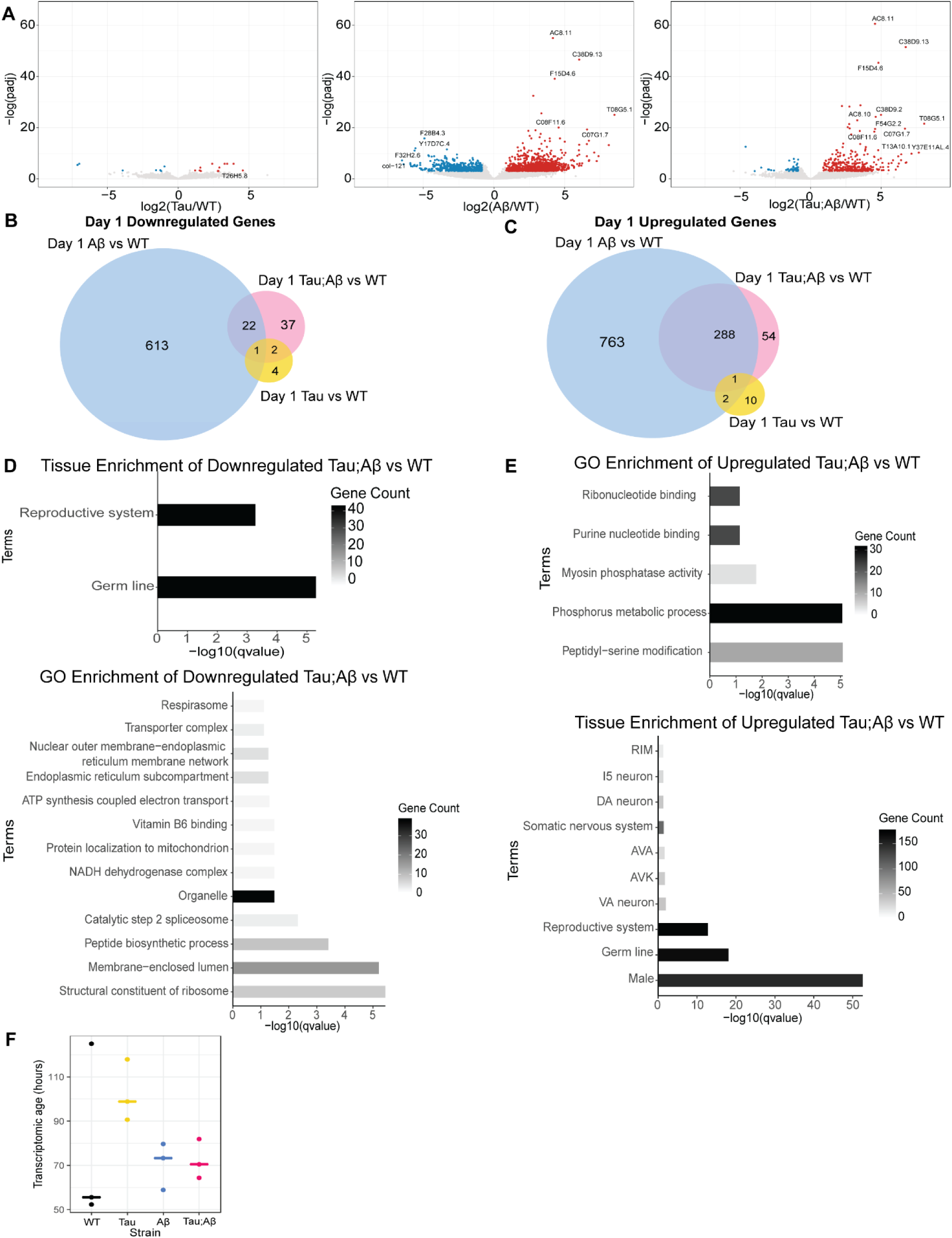
Aβ expression drives transcriptomic changes in the dual Tau;Aβ strain. (A) Volcano plots of fold gene expression change in Tau compared to WT, Aβ compared to WT and Tau;Aβ compared to WT on day 1 of adulthood. (B) Venn diagram of significant genes downregulated and (C) upregulated in Tau, Aβ and Tau;Aβ compared to WT on day 1 of adulthood. (D) Wormbase tissue enrichment and GO enrichment of the genes downregulated in day 1 Tau;Aβ strain compared to WT. (E) Wormbase tissue enrichment and GO enrichment of the genes upregulated in day 1 Tau;Aβ strain compared to WT. (F) Based on transcriptomic aging clock from ^41^, the predicted biological age of Tau, Aβ and Tau;Aβ are increased compared to WT.

